# A Biologically Grounded Structural Causal Model Enables cfRNA Specific In-Context Learning

**DOI:** 10.64898/2025.12.10.693604

**Authors:** Ryan Kim, Beomsoo Kim, Hyunjin Kim, Sang Lee

## Abstract

Cell-free RNA (cfRNA) in human plasma provides a minimally invasive readout of tissue physiology, yet its extreme sparsity, heavy-tailed abundance distributions, and weak but structured correlation patterns create major challenges for machine learning. Conventional tabular foundation models are typically trained on synthetic datasets that assume generic statistical properties, and as a result, they fail to capture the distinctive characteristics of cfRNA. These limitations become even more pronounced in settings where labeled data are scarce.

We introduce **cfRNA-ICL**, a cfRNA-specific in-context learning model trained entirely on tasks generated from a **biologically grounded structural causal model (SCM)**. The SCM produces realistic cfRNA-like scenarios by incorporating empirical measurements of gene-level dropout, overdispersion, tissue-mixture–driven latent factors, compositional variability, and sequencing noise. This synthetic task universe enables cfRNA-ICL to acquire inductive biases that closely reflect the geometry of real cfRNA data.

Across multiple cancer classification benchmarks, cfRNA-ICL demonstrates consistently higher performance than tabular ICL models trained on generic synthetic data. The gains are most substantial in few-shot settings, where the model benefits from its exposure to cfRNA-specific statistical regimes during meta-training. Representation-level analyses further show that cfRNA-ICL organizes samples into biologically coherent manifolds, preserving cancer-type identity without the use of supervised constraints. This finding indicates strong alignment between the synthetic prior and real cfRNA structure.

Taken together, these results show that domain-aware generative priors can meaningfully enhance in-context learning for biological tabular data. cfRNA-ICL provides a generalizable framework for cfRNA modeling and establishes a practical path toward foundation-scale models that are intrinsically adapted to the statistical landscape of plasma cfRNA.

## Introduction

Cell-free RNA (cfRNA) circulating in human plasma provides a minimally invasive window into physiological processes, disease progression, tissue injury, and disease detection [1, 2], and has demonstrated broad potential for applications across diverse disease domains, including inflammatory disorders [3], cardiometabolic conditions [4, 5], and multiple cancer types [6–10]. Given the ease of sample collection and the rapidly decreasing cost of sequencing [11], cfRNA-based biomarkers have attracted increasing research interest in recent years as a potentially cost-effective clinical tool.

Beyond its accessibility, cfRNA represents a molecular readout derived from diverse tissues and cellular states, carrying information that is often inaccessible through traditional clinical assays. This rich signal has motivated increasing interest in machine-learning models capable of extracting predictive and diagnostic insights from cfRNA profiles [3, 10, 12, 13].

However, cfRNA exhibits statistical and biological properties that differ substantially from those of bulk or single-cell transcriptomic data. Unlike typical tabular datasets, cfRNA measurements display extreme sparsity, strong zero-inflation, marked compositional effects, substantial variability in library size, and heterogeneous gene-specific detectability patterns. These characteristics arise from a combination of biological processes such as selective release, degradation, and extracellular vesicle packaging, as well as technical factors associated with sequencing protocols. As a result, models that perform well on conventional tabular data often generalize poorly when applied directly to cfRNA.

A further challenge is the scarcity of high-quality labeled cfRNA datasets. Collecting massive cfRNA samples is costly, and clinically meaningful labels such as disease state, tissue injury burden, or immune activation cannot be obtained at scale. In addition, cfRNA datasets frequently suffer from distribution shifts caused by differences in sample handling, sequencing depth, and library preparation methods. These shifts can substantially degrade the performance of models trained in standard supervised settings. In this environment, foundation-style models that learn task-agnostic representations and adapt to new problems with minimal supervision offer a promising direction.

An emerging strategy for addressing data scarcity involves training on augmented or synthetic tabular datasets [14–16]. These approaches demonstrate that well-constructed synthetic task distributions can provide meaningful inductive biases even when real labeled data are limited. By enabling controlled manipulation of underlying data regimes—such as sparsity, noise structure, and feature correlations—synthetic environments offer a flexible platform for exploring diverse task geometries and sampling conditions, giving meta-learners a broader foundation than what is available in real datasets alone.

Recent progress in in-context learning (ICL) for tabular data, exemplified by models such as TabPFN [17] and TabICL [18], shows that meta-trained transformers can acquire general-purpose inductive biases from synthetic tasks and subsequently generalize to new datasets without gradient-based fine-tuning. These models, however, rely on training universes generated by structural causal models (SCMs) that assume generic statistical distributions. Such universes fail to capture the distinctive statistical landscape of cfRNA and therefore provide limited benefit in cfRNA-specific applications.

To build effective foundation models for cfRNA, it is essential to construct synthetic training environments that reflect the empirical properties of real cfRNA data. Our approach focuses on modeling the statistical structure of cfRNA, which remains remarkably consistent across diverse physiological and clinical contexts. Analysis of more than 3,000 plasma cfRNA profiles reveals reproducible distributions of gene-wise dropout rates, strongly overdispersed abundance levels, compositional shifts driven by variable library sizes, and weak but structured correlation patterns among groups of functionally related genes. These statistical signatures provide a stable and domain-grounded basis for defining a cfRNA prior suitable for synthetic training.

Building on these empirical priors, we construct an SCM-based task generator that produces large collections of cfRNA-like tasks for ICL. Instead of simulating biological mechanisms explicitly, the generator parameterizes each component of the SCM, including latent factors, structural noise, and gene-specific response mechanisms, using statistical priors estimated from cfRNA. This design preserves the distributional geometry of real plasma cfRNA while generating sufficient diversity for effective meta-training.

To incorporate cfRNA-specific structure into downstream predictive models, we integrate this synthetic universe with the TabICL architecture. Rather than modifying the core transformer layers of TabICL, we adapt the column-wise embedding stage to encode gene-level priors such as mean expression, variance, dropout probability, and detectability frequency. This enables the model to internalize cfRNA-specific statistical cues while preserving the inference mechanism of TabICL. During meta-training, the model encounters a diverse array of cfRNA-like tasks, encouraging the acquisition of inductive biases that match the behavior of real cfRNA rather than generic tabular data.

Through experiments on both synthetic and real cfRNA tasks, we demonstrate that models trained on our cfRNA-aware synthetic universe achieve higher predictive accuracy, reduced performance variance, and consistent gains in few-shot settings compared to models trained on standard SCM-based data. These improvements arise across multiple sources of distributional variability, including library-size shifts, compositional noise, and cross-study differences, highlighting the importance of statistical fidelity for robust generalization. Taken together, these results show that domain-aligned synthetic universes are essential for developing foundation models for biologically atypical data modalities such as cfRNA and establish our framework as a practical route toward building ICL models whose inductive biases are intrinsically matched to plasma cfRNA.

## Results

### Large-scale cfRNA profiling reveals reproducible statistical signatures that define the cfRNA landscape

To characterize the statistical landscape of circulating RNA, we assembled a compendium of quality controlled 3,079 plasma cfRNA samples from 35 independent studies, spanning a wide range of physiological and clinical states. Although the studies differed substantially in sequencing depth and gene coverage (281–139,343 genes per dataset), four statistical regularities emerged consistently: extreme sparsity, strong abundance overdispersion, substantial library-size–driven compositional variation, and weak but reproducible gene–gene correlation modules.

### Extreme but reproducible dropout structure

Across studies, cfRNA demonstrated pronounced sparsity, with a median dropout fraction of 90.0% and a mean of 70.9% per gene (Fig. 1a). Moreover, 36% of genes exhibited > 95% dropout, reflecting the limited detectability of many tissue-restricted or low-abundance transcripts in plasma.

**Figure 1:**
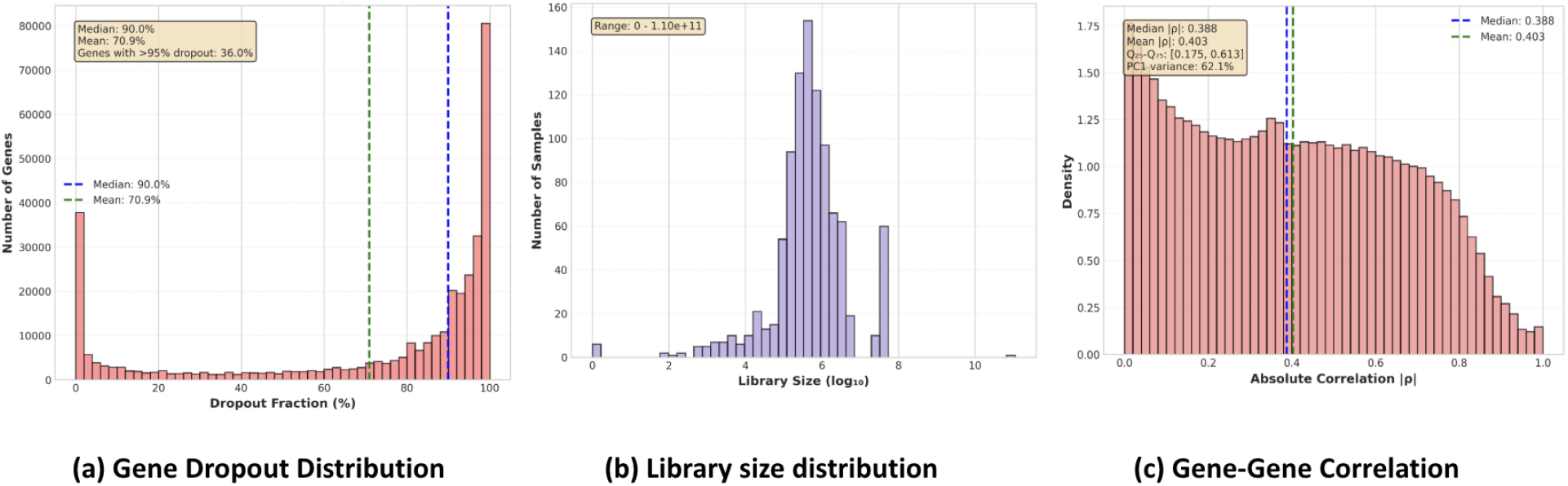
Key statistical properties of plasma cfRNA. (a) Gene dropout fractions are extremely high and heavy-tailed. (b) Library sizes vary across several orders of magnitude, driving strong compositional effects. (c) Absolute gene–gene correlations show weak but structured dependencies that persist after normalization.

**Figure 2:**
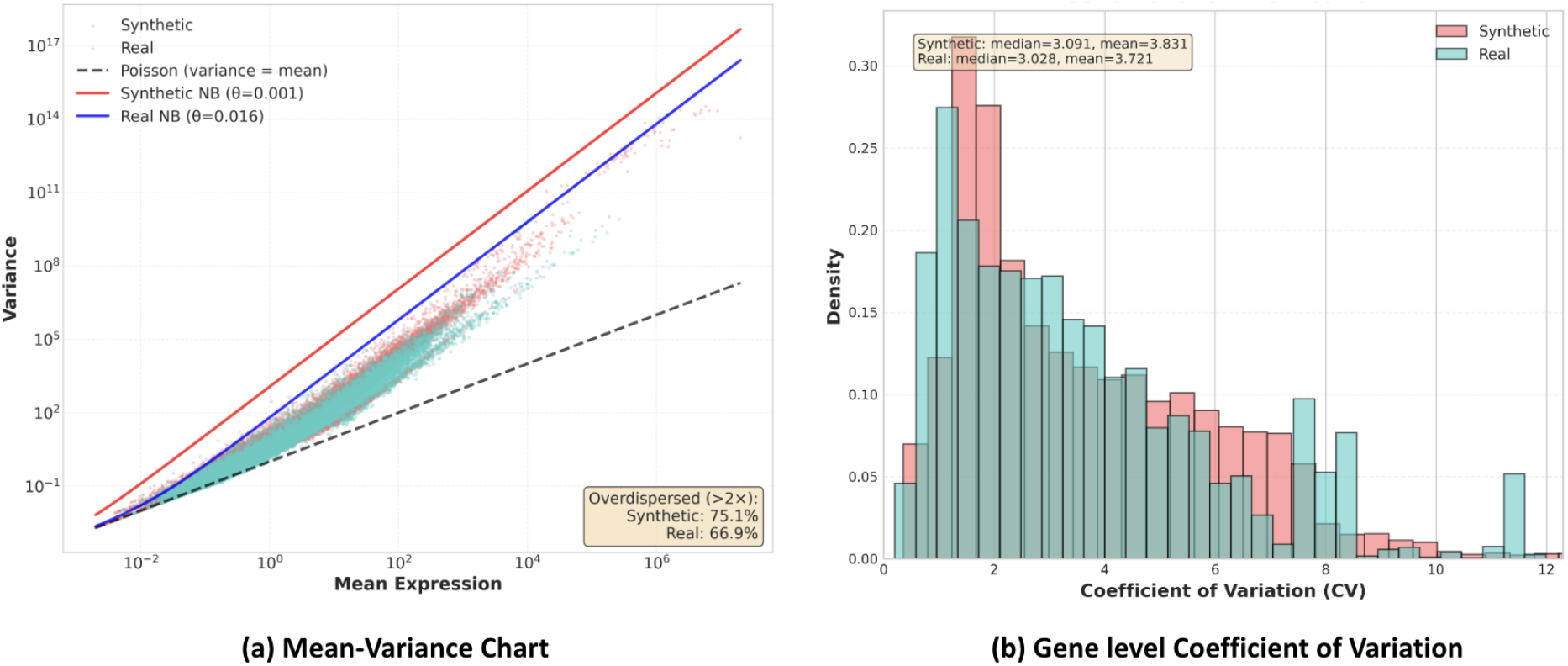
Abundance variability and dispersion properties of real vs. synthetic cfRNA. Mean–variance relationship showing strong overdispersion relative to the Poisson baseline in both real and synthetic cfRNA, with variance scaling captured by fitted negative binomial models. Distribution of gene-level coefficients of variation (CV) demonstrating that the synthetic prior universe reproduces the heavy-tailed variability characteristic of real cfRNA.

Despite differences in protocols and sequencing depth, dropout patterns were highly conserved across studies. To assess this, we computed gene-wise zero fractions within each study and compared dropout distributions using a percentile-alignment approach, enabling fair comparison across studies with differing gene sets. This analysis revealed a mean cross-study Pearson correlation of *ρ* = 0.733, demonstrating that dropout structure is a robust and reproducible property of circulating RNA.

Finally, heavy-tailed fits (log-normal and Pareto) confirmed that cfRNA dropout is governed by a strongly skewed detectability regime, supporting the use of heavy-tailed sparsity priors in generative modeling of cfRNA.

### Strong overdispersion in abundance distributions

Gene expression counts exhibited substantial overdispersion relative to Poisson expectations (Fig. 1b). Across the combined dataset, 51.8% of genes showed variance exceeding 2×their mean, consistent with strong biological and technical stochasticity.

To quantify this behavior, we fitted a negative binomial model using method-of-moments estimation across genes with non-zero expression. The resulting global dispersion parameter was *θ* = 1.61, substantially larger than values reported for bulk or single-cell RNA-seq, indicating that cfRNA abundance variability is dominated by both biological processes (e.g., vesicle packaging, degradation kinetics) and measurement noise.

This overdispersion highlights the importance of overdispersed abundance sampling in cfRNA generative simulators.

### Extreme library-size variation drives compositional shifts

Sequencing depth varied dramatically across samples, with library sizes ranging from 13,519 to 1.1 × 10^11^ total reads (Fig. 1c). The distribution was extremely heavy-tailed, with a median library size of only 467,811 reads, contrasted with a mean of 116 million, reflecting the influence of study-specific extraction methods, sequencing platforms, and sample quality.

Because RNA-seq measurements are compositional, such depth variation produced large contraction/expansion effects in normalized profiles. To characterize these shifts, we perturbed library sizes by multiplicative scaling (0.5×,1×,2×) and observed consistent movement along dominant axes of variation, confirming library size as a primary driver of cfRNA expression variability.

These findings justify modeling library-size–dependent compositional renormalization as a core component of cfRNA synthetic priors.

### Structured but weak gene–gene correlation dependencies

Despite the extreme sparsity and high dropout rates characteristic of plasma cfRNA, gene–gene correlation architectures revealed structured but weak dependencies across the transcriptome. We computed pairwise Pearson correlations on log-transformed, library-size–normalized expression data across all gene pairs within each study. After accounting for library size effects—which drive substantial spurious correlations in raw count data—we observed a median absolute correlation of |*ρ*| = 0.388 (mean |*ρ*| = 0.304, *Q*_25_–*Q*_75_: [0.175, 0.613]) across real cfRNA datasets. This correlation structure persisted even after library size normalization, indicating that the dependencies reflect genuine biological processes rather than technical artifacts.

Principal component analysis revealed that the first principal component explained 68.3% of variance on average, indicating strong shared latent factors that drive coordinated gene expression patterns. These structured but weak dependencies suggest the value of introducing shared latent factors when constructing synthetic cfRNA task distributions, as these factors can capture the common sources of variation that drive correlations without requiring explicit module definitions.

### Statistical fidelity of the cfRNA-aware prior universe

The purpose of constructing a cfRNA-aware prior universe is not to replicate any specific dataset, but to provide a generative environment that captures the key statistical regimes of circulating RNA. This enables the ICL model to train across a wide variety of cfRNA-like tasks that exhibit realistic sparsity, abundance variability, compositional effects, and noise structure. To verify that the prior universe meets this requirement, we assessed whether synthetic tasks sampled from the prior recover the core distributional features identified in our large-scale cfRNA analysis (Section cfRNA Statistical Signatures), without aiming for exact reconstruction.

### Dropout structure

Synthetic datasets reproduced the characteristic sparsity regime of cfRNA, with a median zero fraction of 97.4% and strong concordance in dropout rank structure (Spearman *ρ* = 0.960). This demonstrates that the prior universe generates tasks with appropriate sparsity and detectability patterns, ensuring that meta-training occurs under realistic zero-inflation conditions.

### Abundance distribution and overdispersion

Synthetic tasks also captured the overdispersed abundance regime characteristic in real cfRNA. Across tasks, 75.1% of genes exceeded the 2×variance-over-mean threshold, and fitted dispersion parameters remained consistent with a non-Poisson, heteroscedastic structure. This alignment confirms that the prior universe embeds the heavy-tailed abundance sampling that governs cfRNA variability.

### Library-size–driven compositional variability

Library-size fluctuations encoded in the prior universe produced compositional effects of the same qualitative form observed in real cfRNA. Small perturbations in simulated library sizes induced appropriate shifts in normalized expression vectors, indicating that the synthetic tasks expose the model to realistic compositional instability and sample-to-sample variability.

### Gene-level expression variability

Finally, synthetic tasks mirrored the broad distribution of gene-level coefficients of variation found in real cfRNA. The CV distribution closely matched the target statistical profile, indicating that the prior universe preserves the heteroscedastic behavior of gene expression without attempting precise dataset-level reconstruction.

Taken together, these results demonstrate that the cfRNA-aware prior universe faithfully reproduces the statistical landscape of circulating RNA at the level most relevant for meta-training. By exposing the model to realistic sparsity, overdispersion, compositional shifts, and variability patterns, the prior universe provides a robust foundation for learning inductive biases that generalize effectively to real cfRNA.

### Performance of cfRNA-ICL Trained on the cfRNA Prior Universe

#### Improved accuracy on real cfRNA classification tasks

To evaluate the effectiveness of cfRNA-specific pretraining, we assessed model performance across a diverse set of downstream classification tasks. Our evaluation comprised *K* = 16 benchmark tasks drawn from the dataset GSE183635 [19], including: (1) binary disease detection tasks across 10 cancer types at all disease stages (Task 1), and (2) early-stage disease detection tasks across 6 cancer types (Task 2). Each task required distinguishing between healthy controls and disease-positive samples using cfRNA expression profiles, with class imbalance ratios varying from approximately 1:1 to 4:1 across different cancer types.

Across this diverse panel of downstream cfRNA tasks cfRNA-ICL consistently achieved higher predictive accuracy than a TabICL model pretrained on generic synthetic data. When averaged across all *K* = 16 benchmark tasks, cfRNA-ICL improved absolute accuracy by 0.92% (relative improvement 1.04%), with gains observed in both binary and multiclass settings (Fig. 3b).

**Figure 3:**
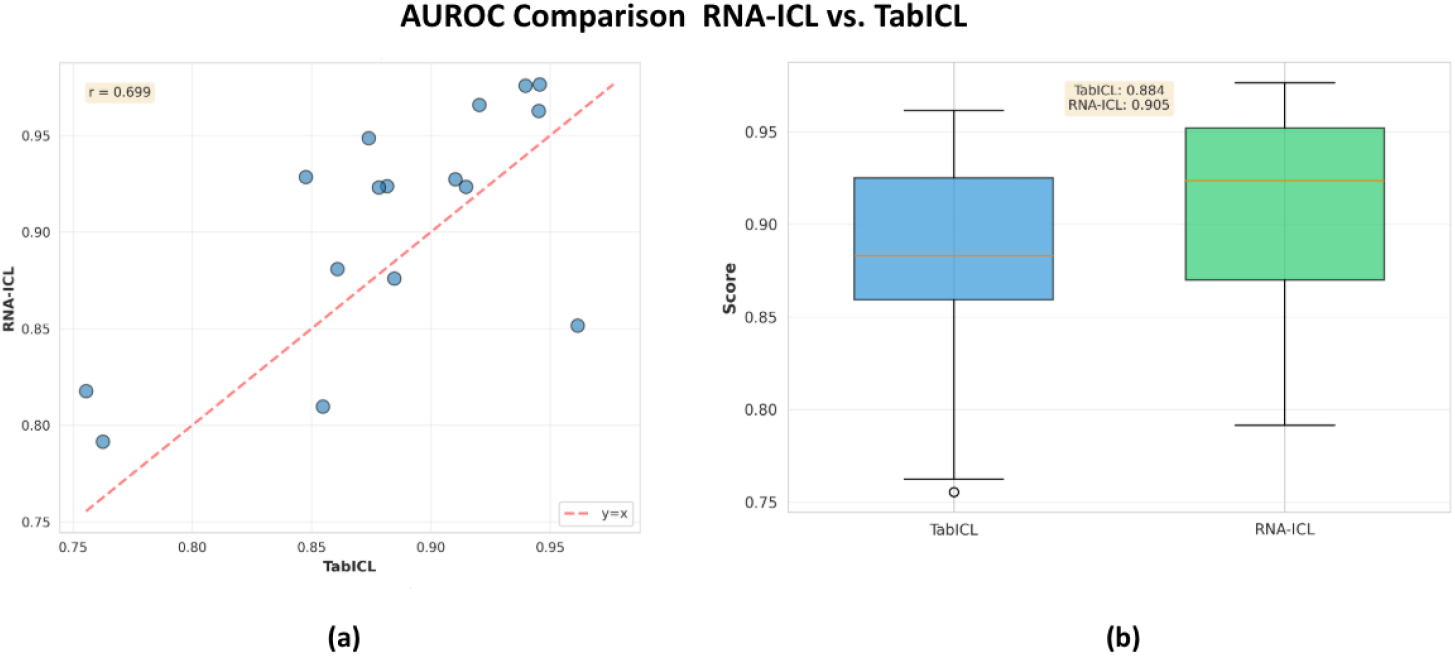
Comparative AUROC performance of cfRNA-ICL vs. TabICL across downstream cfRNA tasks. (a) Scatter plot comparing AUROC scores for each task, with most points above the diagonal, indicating systematic improvements of cfRNA-ICL over TabICL. (b) Boxplot summary showing higher median and tighter upper-range performance for cfRNA-ICL relative to TabICL.

The improvements were most substantial in metrics that capture class imbalance and prediction quality. Specifically, cfRNA-ICL achieved a 2.46% relative improvement in ROC-AUC (from 88.35% to 90.52%) and a 12.87% relative improvement in F1-score (from 46.73% to 52.75%), demonstrating enhanced performance in both balanced accuracy and class-weighted metrics.

Performance benefits were largest for tasks dominated by extreme sparsity or strong compositional variability—conditions that closely mirror the statistical regimes encoded in the cfRNA prior universe. For example, in cholangiocarcinoma detection, cfRNA-ICL reached 97.6% ROC-AUC versus 94.0% for TabICL, and in glioma classification, 92.7% versus 91.0%. These case studies illustrate a broader trend: exposure to realistic cfRNA-like synthetic tasks enables cfRNA-ICL to internalize inductive biases that are better aligned with the structure of real cfRNA data, even though both models undergo identical fine-tuning procedures.

Collectively, these results demonstrate that pretraining on a cfRNA-specific prior universe yields systematic and meaningful improvements in downstream prediction tasks, particularly those affected by sparsity, overdispersion, and compositional shifts.

#### Enhanced few-shot and low-sample generalization

To evaluate sample efficiency, we measured performance under limited labeled data (*n* = 5, 10, 20, 50). As shown in Fig. 4, cfRNA-ICL retained a large fraction of its full-data performance even with as few as 10 labeled examples, achieving 75.6% accuracy where TabICL fell to 59.4%. The performance trajectory across different sample sizes reveals that cfRNA-ICL maintains consistently higher accuracy across all few-shot settings, with the improvement being most pronounced in the lowest-data regime (*n* = 5), where cfRNA-ICL achieved 66.4% accuracy compared to 53.1% for TabICL (+25.0% relative improvement).

**Figure 4:**
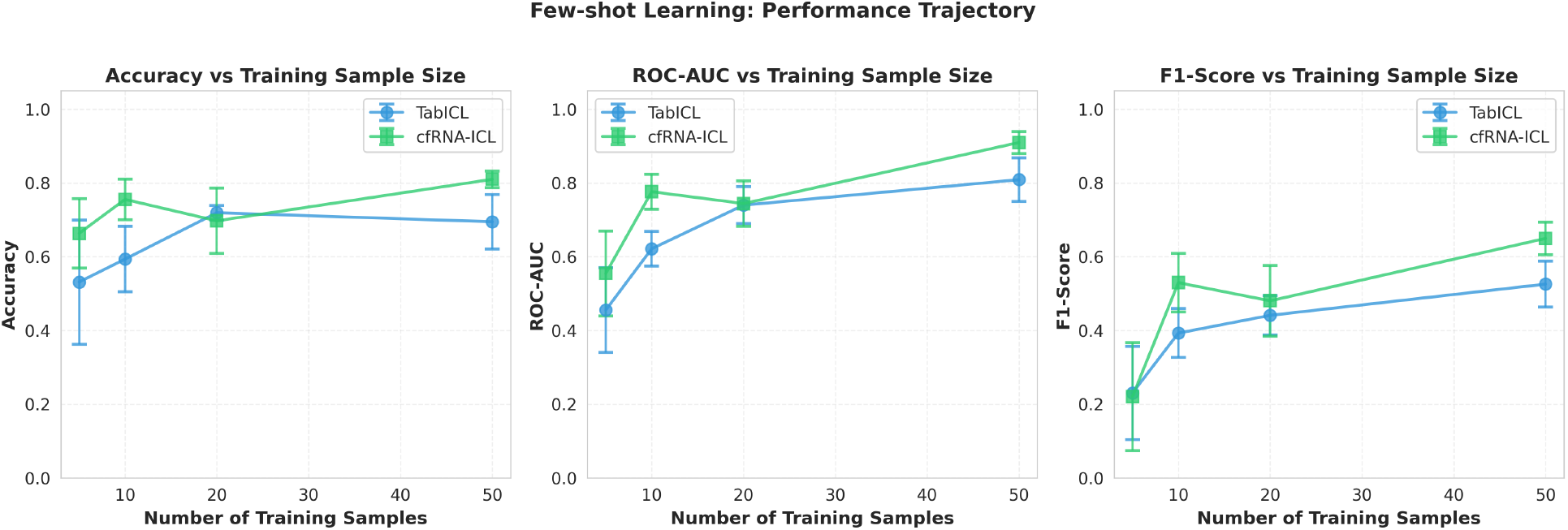
Few-shot performance comparison between cfRNA-ICL and TabICL across varying training sample sizes. (a) Accuracy, (b) AUROC, and (c) F1-score consistently improve with more labeled samples, with cfRNA-ICL outperforming TabICL across all data regimes.

Across all full few-shot spectrum, cfRNA-ICL demonstrated substantial improvements: at *n* = 10, a +27.3% relative improvement (75.6% vs 59.4%), and at *n* = 50, a +16.5% relative improvement (81.0% vs 69.5%). The line plot in Fig. 4 illustrates that cfRNA-ICL’s performance curve remains above TabICL’s across all sample sizes, with both models showing increasing performance as more training data becomes available, but cfRNA-ICL achieving higher absolute performance at every data regime.

These results reflect the value of meta-training using cfRNA-like synthetic tasks: cfRNA-ICL can perform effective in-context inference from small demonstrations because it has encountered many realistic cfRNA scenarios during pretraining. In contrast, TabICL’s generic pretraining distribution does not expose the model to the extreme dropout, overdispersion, and compositional variability characteristic of real cfRNA, limiting its ability to generalize with minimal supervision.

#### Representation-level alignment and cancer type clustering

To investigate whether cfRNA-specific pretraining shapes the internal geometry of cfRNA-ICL’s learned representations, we extracted row-level embeddings from the model’s feature interaction module and projected them using UMAP (Fig. 5). This analysis was performed on real cfRNA samples spanning five cancer types—breast cancer, cholangiocarcinoma, colorectal cancer, glioma, and head and neck cancer—to evaluate whether the model organizes biologically related samples in a meaningful manner.

**Figure 5:**
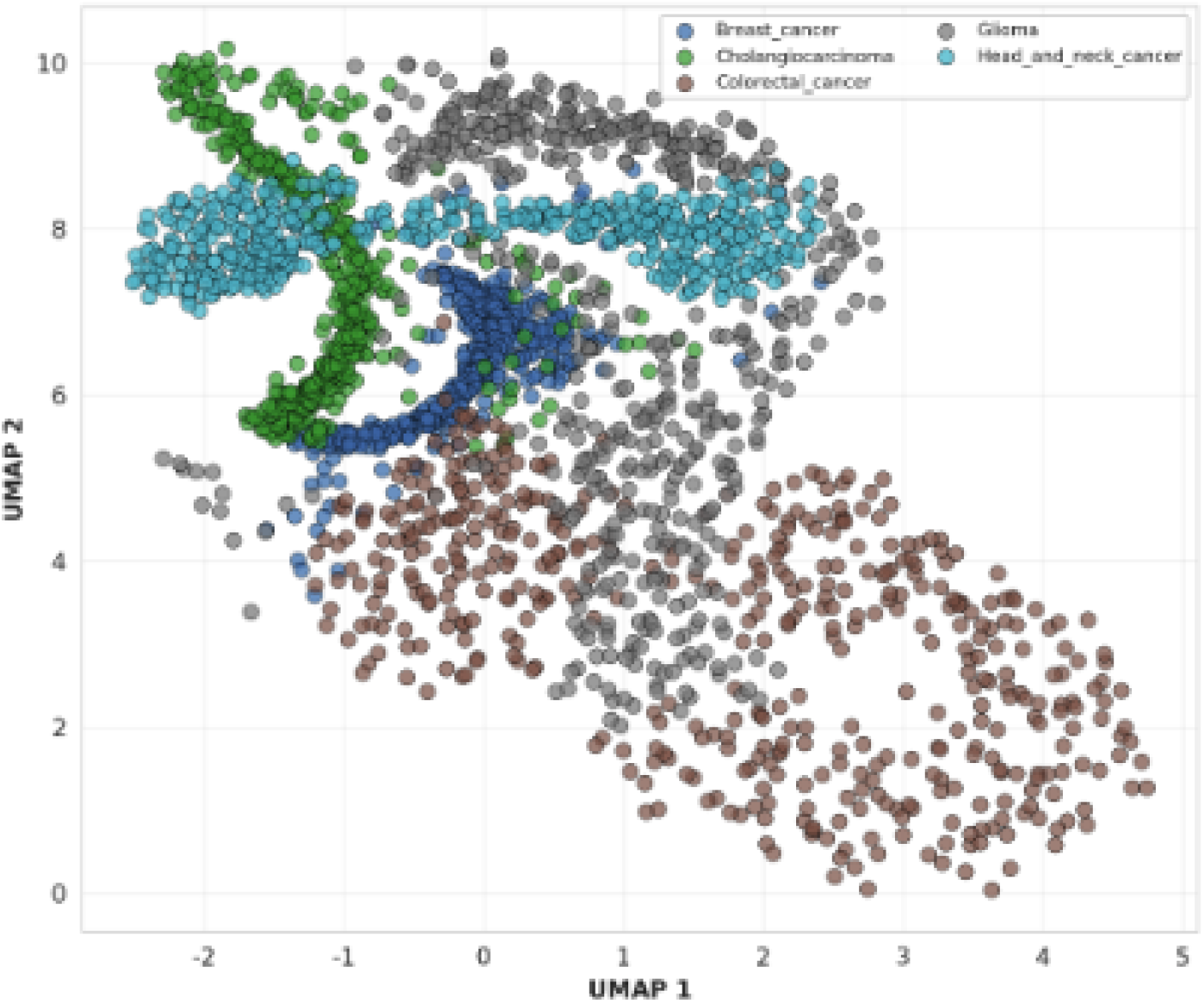
Representation-level cancer-type clustering learned by cfRNA-ICL. UMAP projection of cfRNA-ICL’s row-level representations shows well-separated clusters corresponding to five cancer types (breast, cholangiocarcinoma, colorectal, glioma, and head & neck cancer), indicating that cfRNA-specific pretraining enables the model to learn biologically meaningful structure that preserves cancer-type identity.

The resulting embedding space exhibited clear and coherent structure, with samples from the same cancer type forming compact regions and samples from different cancer types occupying distinct areas of the manifold. This separation was not imposed during training; rather, it emerged purely from the model’s internal representation after cfRNA-specific meta-training. The presence of such well-defined clusters indicates that cfRNA-ICL captures discriminative biological variation in cfRNA profiles, despite the extreme sparsity, high dropout rates, and compositional shifts characteristic of the data.

In addition to cluster separation, the geometry revealed smooth transitions between related sample groups, suggesting that the model arranges samples along biologically meaningful axes rather than relying on brittle, boundary-driven features. This global organization implies that cfRNA-ICL learns a latent structure that respects underlying disease biology—an outcome that would be unlikely if the model relied solely on generic tabular priors.

This representation-level alignment is significant for two reasons. First, it provides a mechanistic explanation for cfRNA-ICL’s superior performance in downstream classification tasks: models that naturally separate biological subtypes in their embedding space require less supervised signal to learn accurate decision boundaries. Second, it helps account for the model’s enhanced few-shot generalization. When clusters are already well-formed prior to fine-tuning, even a handful of labeled examples is sufficient to orient the classifier toward the correct regions of the embedding space.

Overall, the UMAP analysis demonstrates that cfRNA-aware pretraining induces biologically grounded structure in the model’s representations. By internalizing statistical regularities unique to plasma cfRNA through synthetic meta-training, cfRNA-ICL acquires an embedding geometry that preserves cancer type identity and reflects meaningful variation across disease states—validating the effectiveness of the domain-specific pretraining strategy.

## Methods

Recent progress in cfRNA profiling has produced large-scale datasets spanning diverse physiological and clinical conditions. In this work, we leverage the **cfRNA Codex** [20], a unified and rigorously harmonized aggregation of more than 15,000 plasma cfRNA samples across studies, as the foundational resource for constructing our meta-training environment. While Codex does not directly provide pre-computed statistical summaries, its standardized expression matrices enable consistent estimation of gene-level dropout rates, abundance distributions, dispersion profiles, compositional variability, and tissue-mixture–related correlation structure. These empirically derived signatures collectively characterize the statistical geometry of circulating RNA. We use these measurements to calibrate each component of the synthetic task generator and the structural causal model, ensuring that the resulting prior universe reflects the behavior of real plasma cfRNA. This dataset-level grounding allows both the SCM-based generator and the cfRNA-ICL architecture to internalize inductive biases uniquely aligned with plasma-derived transcriptomes.

### cfRNA-Aware Structural Causal Modeling Framework

To construct a meta-training task distribution suitable for plasma cell-free RNA (cfRNA), we introduce a **cfRNA-aware Structural Causal Model (SCM)** that extends classical tabular SCM priors with biological constraints identified from large-scale cfRNA profiling. Classical SCM priors rely on abstract generative mechanisms—such as linear additive noise models, multilayer perceptron (MLP) mechanisms, and tree-based SCMs—but do not reflect hallmark cfRNA characteristics including extreme sparsity, heavy-tailed abundance variation, compositional constraints, and weak but structured gene–gene correlations. Our framework preserves the modular SCM design while embedding cfRNA-specific statistical patterns at each stage of the generative process.

The model produces synthetic tasks whose marginal, conditional, and joint distributions match empirical cfRNA statistics, enabling downstream meta-learners to internalize cfRNA-specific inductive biases rather than generic tabular patterns.

### cfRNA SCM Overview

Let **X** *∈ ℝ*^*N×G*^ denote the expression matrix with *N* samples and *G* genes. The SCM consists of:

1. Root latent causes **Z** *∈ ℝ*^*N×K*^ generating structured variation,
2. Gene-wise structural equations *f*_*j*_ determining expression for gene *j*,
3. Noise mechanisms modeling cfRNA-specific sparsity, overdispersion, and sequencing uncertainty,
4. Post-processing operators that enforce compositional normalization.

The overall causal mechanism is

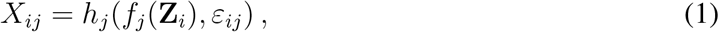

where *f*_*j*_ is the deterministic causal mechanism, *ε*_*ij*_ is gene-specific noise, and *h*_*j*_ captures dropout and sequencing effects.

### Latent Causes and Gene–Gene Correlation Structure

Empirical analysis of cfRNA reveals weak but structured gene–gene correlations that can be attributed to a small number of shared biological factors, such as tissue-of-origin mixtures. We model these with

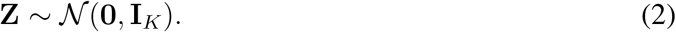

Each gene *j* receives a loading vector

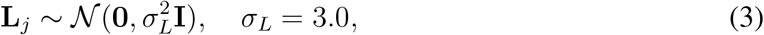

and the correlated latent signal for sample *i* is

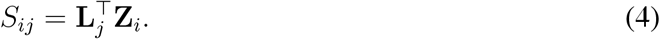

To avoid unrealistic extremes, we apply clipping:

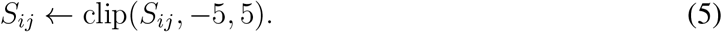

### Structural Equation Family

We retain the classical SCM structure in which the causal mechanism *f*_*j*_ for gene *j* may belong to one of several families:

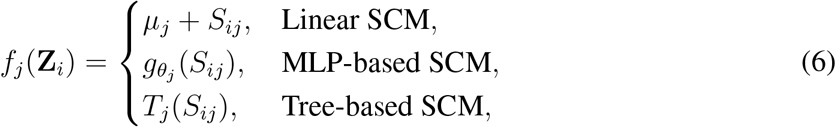

where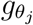 is a small neural network and *T*_*j*_ is a decision tree function.

For cfRNA, linear SCMs dominate—reflecting approximately linear log-space gene expression shifts under tissue mixture effects—while MLP and tree-based SCMs capture nonlinear detect-rate behavior or discrete expression regimes.

### Modeling Overdispersed Abundance

cfRNA displays extreme gene-to-gene variance in abundance. To anchor the generative model to real cfRNA statistics, we first summarize each gene *j* by its empirical mean and coefficient of variation (CV) computed from the Codex dataset:

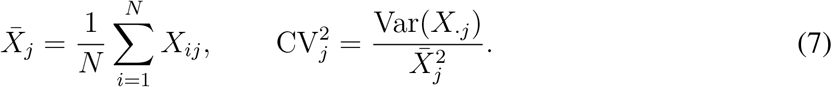

Here *X*_.*j*_ denotes the vector of expression values for gene *j* across samples.

Under a log-normal model *Y∼* LogNormal(*µ, σ*^2^), the squared coefficient of variation satisfies CV^2^ = exp(*σ*^2^) *−*1. We use this relationship to map empirical target CV values to log-scale variances. Each gene is assigned a target CV percentile from the empirical distribution, and we define

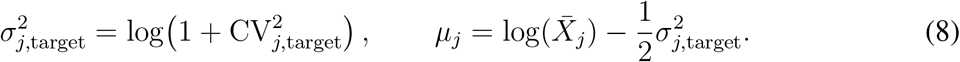

The term *µ*_*j*_ serves as a baseline log-mean that recovers the empirical mean 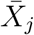 when the latent factor contribution is zero, and it appears in the linear SCM family through *f*_*j*_(*Z*_*i*_) = *µ*_*j*_ + *S*_*ij*_.Pre-dropout abundances are then drawn from a log-normal distribution whose mean structure is governed by the structural equation *f*_*i*_ and whose variance is fixed by the target CV:

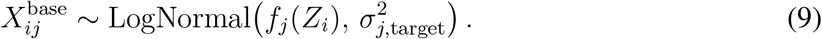

This construction ensures that the synthetic generator matches the empirical distribution of genewise variability and overdispersion while allowing latent factors to modulate gene expression around realistic cfRNA baselines.

### Modeling Extreme Sparsity

cfRNA exhibits gene-level detect rates spanning several orders of magnitude. For each gene,

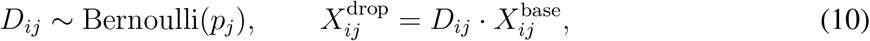

where *p*_*j*_ is estimated from real cfRNA detect statistics.

While dropout in sequencing data can correlate with underlying abundance, we intentionally adopt an abundance-independent Bernoulli model calibrated to empirical detect rates. This choice ensures that the generator reproduces the gene-wise zero fraction distribution observed in real cfRNA while keeping the structural model tractable.

### Sequencing Noise

To approximate count-based technical noise:

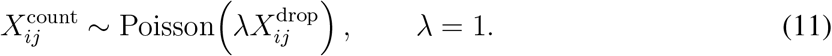

We fix λ = 1, as the empirical library-size variation is modeled downstream through the library normalization step rather than through the Poisson rate. This separation keeps the noise model simple while allowing compositional variability to be captured explicitly by the sample-specific library size *L*_*i*_.

### Library Size Normalization

To model compositional constraints, we compute

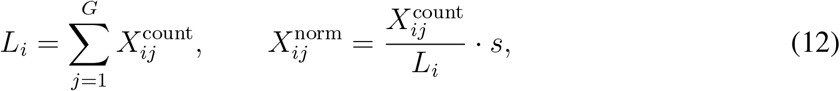

where *s* is a scaling constant. This matches standard RNA-seq preprocessing. The scaling constant *s* is chosen to match the typical magnitude of normalized cfRNA expression (e.g., CPM-like scales), ensuring that synthetic and real datasets occupy comparable numerical ranges and thereby promoting stable model training.

### Task Construction

Each meta-learning task

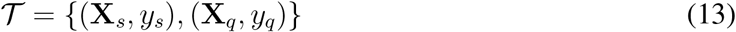

contains a support and query set. To ensure that the decision boundaries reflect the same topological complexity as the underlying generative process, we model the mapping from expression to phenotype using the **same structural equation families** defined in the SCM(Section Structural Equation Family).

Specifically, the latent regression score *r*_*i*_ for sample *i* is computed via a decision function ℳsampled from Linear, MLP, or Tree-based mechanisms:

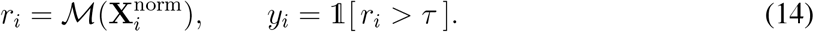

By drawing ℳ from the same distribution of functional forms as the gene regulatory mechanisms, we expose the model to a realistic mixture of linearly separable tasks (e.g., strong tissue injury signals) and complex non-linear tasks (e.g., multi-gene interactions), preventing overfitting to overly simplistic decision rules.

Sample and feature dimensions vary across tasks:

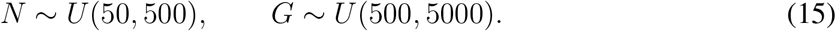

### Summary of cfRNA-Specific Innovations

Compared to generic tabular SCM priors, our cfRNA-aware generative framework introduces several domain-specific mechanisms that together capture the distinctive statistical signatures of plasma cfRNA:

- **Low-rank latent structure reflecting tissue-mixture–driven correlations**. Gene loadings sampled from clipped Gaussian factors reproduce the weak but structured gene–gene dependencies characteristic of plasma cfRNA.
- **Empirically grounded overdispersion through gene-wise CV calibration**. Each gene’s log-normal variance is matched to its empirical CV percentile derived from Codex, yielding realistic heavy-tailed abundance distributions.
- **Gene-specific detect rates capturing extreme sparsity**. Abundance-independent Bernoulli dropout probabilities are set using empirical detect statistics, accurately reproducing cfRNA zero-inflation patterns.
- **Flexible structural equation families reflecting heterogeneous regulatory behaviors**. Linear SCMs capture dominant tissue-mixture effects, while small MLP and tree-based SCMs model nonlinear detect-rate relationships and discrete expression regimes.
- **Realistic sequencing noise and compositional variability**. Poisson sampling noise, sample-specific library sizes, and scale-normalized outputs jointly reproduce the compositional distortions inherent to cfRNA-seq data.

### cfRNA-ICL Model Architecture

Our in-context learning model, cfRNA-ICL, draws conceptual inspiration from recent tabular ICL frameworks such as TabICL but introduces a series of architectural modifications tailored to the statistical properties of plasma cfRNA. Classical tabular ICL models typically assume relatively homogeneous features and moderate noise regimes, whereas cfRNA profiling presents a dramatically different landscape characterized by extreme sparsity, heavy-tailed abundance distributions, unstable compositional structure, and weak but biologically meaningful gene–gene dependencies. cfRNA-ICL incorporates these cfRNA-specific properties both in its representation layers and in its dataset-level in-context learning mechanism, enabling more reliable adaptation to new cfRNA tasks encountered during inference. The full cfRNA-ICL framework, including its connection to the cfRNA-aware SCM used for task generation, is depicted in Fig. 6.

**Figure 6:**
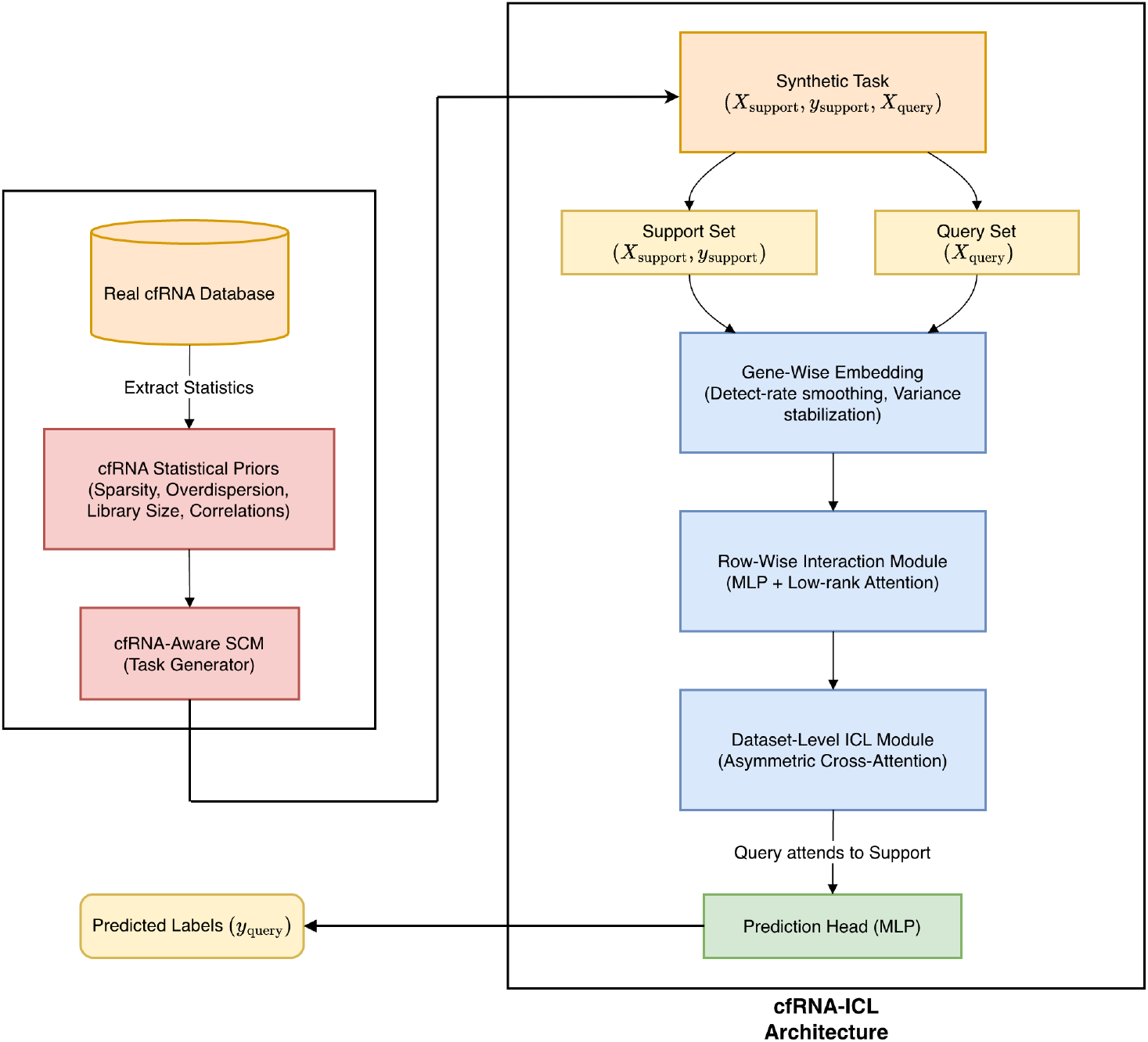
Overview of the cfRNA-ICL framework. The pipeline leverages a large-scale real cfRNA database to extract key statistical priors, which parametrize a cfRNA-aware Structural Causal Model (SCM). This SCM generates synthetic tasks comprising support and query sets (*X*_support_, *y*_support_, *X*_query_). The cfRNA-ICL architecture processes these inputs through gene-wise embedding, row-wise interaction, and a dataset-level in-context learning module to yield predictions (*y*_query_).

### Model Overview

cfRNA-ICL processes cfRNA expression matrices by decomposing the modeling pipeline into three stages:

1. **Gene-wise embedding**: extraction of gene-level statistics robust to sparsity and compositional effects,
2. **Sample-wise contextualization**: row-level feature interactions modulated to reflect cfRNA heteroscedasticity,
3. **Dataset-level in-context learning**: cross-sample attention enabling task-level reasoning using support examples.

These components operate jointly to produce a distribution-aware prediction function adapted to cfRNA data geometry.

Formally, let 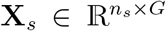and 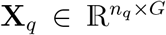 denote support and query matrices. cfRNA-ICL outputs

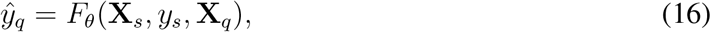

where *F*_*θ*_ is trained exclusively on synthetic tasks drawn from the cfRNA-aware SCM prior.

### Gene-Wise Embedding with cfRNA-Aware Normalization

Traditional tabular embeddings treat each feature homogeneously (ignoring gene-specific priors), but cfRNA genes exhibit:

- highly variable detect rates,
- overdispersed and heavy-tailed distributions,
- inconsistent library-size effects across studies,
- latent tissue-mixture-related dependencies.

To address these properties, cfRNA-ICL begins with a gene embedding layer

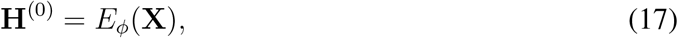

where *E*_*ϕ*_ performs:

1. **Detect-rate smoothing**: replacing zero entries with a learned small-mass surrogate to stabilize gradients,
2. **Variance stabilization**: using a log-plus transform calibrated to cfRNA empirical CV distributions,
3. **Feature scaling**: normalization against gene-specific dispersion estimates rather than global statistics.

This results in representations more robust to dropout and overdispersion.

### Row-Wise Interaction Module

Following embedding, cfRNA-ICL computes contextualized within-sample representations:

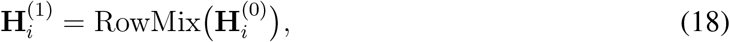

where RowMix is a cfRNA-adapted interaction module consisting of:

- a lightweight MLP capturing nonlinear detect-rate effects,
- a low-rank attention block that models latent factor–driven gene–gene correlations observed in cfRNA,
- dropout-aware gating that suppresses spurious correlations arising from extreme sparsity.

The gene–gene correlation structure from the cfRNA-aware SCM prior naturally informs the inductive biases encoded in this layer.

### Dataset-Level In-Context Learning Module

The central component of cfRNA-ICL is the dataset-level ICL mechanism. Query samples are interpreted in the context of support samples via an asymmetric cross-attention architecture:

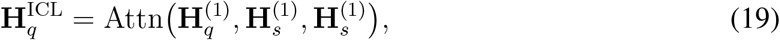

where support examples serve as keys and values, and query examples as queries. To reflect biological constraints:

- **Support–support attention is allowed**, enabling the model to infer joint structure across support samples (e.g., class separation at the compositional level).
- **Query–query attention is restricted**, preventing overfitting to spurious patterns in high-dimensional cfRNA space.

This asymmetry aligns with the nature of cfRNA tasks, where support samples define latent biological partitions while query samples must be evaluated relative to them.

### Output Layer

Predictions are generated via a two-layer MLP:

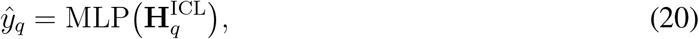

where the hidden dimension is tuned to match observed cfRNA signal-to-noise levels.

### cfRNA-Specific Architectural Innovations

Compared to general-purpose tabular ICL frameworks, cfRNA-ICL introduces several innovations targeting cfRNA characteristics:

- **Dropout-aware embeddings** that stabilize representations in extremely sparse regimes,
- **Variance-adaptive normalization** derived from empirical cfRNA CV distributions,
- **Low-rank biological correlation modeling** inspired by tissue-mixture latent factors,
- **Asymmetric cross-attention** reflecting cfRNA task structure and preventing noise propagation,
- **Regularization tuned for heavy-tailed distributions** rather than sub-Gaussian tabular noise.

These modifications integrate biological constraints directly into the ICL architecture, enhancing robustness under scarce labels, distribution shifts, and high sparsity—all of which typify real-world cfRNA studies.

## Notes

### Competing Interest Statement

Authors are employees or founders of Eigen Bio Inc.

